# Optimizing the Electric Field Strength in Multiple Targets for Multichannel Transcranial Electric Stimulation

**DOI:** 10.1101/2020.05.27.118422

**Authors:** Guilherme B. Saturnino, Kristoffer H. Madsen, Axel Thielscher

## Abstract

**Objective:** Most approaches to optimize the electric field pattern generated by multichannel Transcranial Electric Stimulation (TES) require the definition of a preferred direction of the electric field in the target region(s). However, this requires knowledge about how the neural effects depend on the field direction, which is not always available. Thus, it can be preferential to optimize the field strength in the target(s), irrespective of the field direction. However, this results in a more complex optimization problem.

**Approach:** We introduce and validate a novel optimization algorithm that maximizes focality while controlling the electric field strength in the target to maintain a defined value. It obeys the safety constraints, allows limiting the number of active electrodes and allows also for multi-target optimization.

**Main Results:** The optimization algorithm outperformed naïve search approaches in both quality of the solution and computational efficiency. Using the amygdala as test case, we show that it allows for reaching a reasonable trade-off between focality and field strength in the target. In contrast, simply maximizing the field strength in the target results in far more extended fields. In addition, by maintaining the pre-defined field strengths in the targets, the new algorithm allows for a balanced stimulation of two or more regions.

**Significance:** The novel algorithm can be used to automatically obtain individualized, optimal montages for targeting regions without the need to define preferential directions. It will automatically select the field direction that achieves the desired field strength in the target(s) with the most focal stimulation pattern.

## 1. Introduction

Transcranial electric stimulation (TES) methods inject weak direct or alternating currents via scalp electrodes in order to create an electric field in the brain that modulates neural activity [1]. In order to improve the robustness of the stimulation outcome and to be able to causally relate the behavioral stimulation effects with the modulation of specific brain areas, it is important to limit the electric field region to one or more regions of interest. However, as the electric field is shaped by the individual head anatomy [2], targeting TES electric fields is not a trivial task. Therefore, multiple optimization approaches have been proposed in order to automatically plan multichannel TES interventions [3–9].

Most TES optimization methods aim to maximize, control or approximate projections of the electric field in a specific direction in the target region, rather than the absolute field strength (or norm) irrespective of direction. This can be a good choice for many cortical targets, as it is thought that the physiological TES effects are direction dependent, such that an electric field pointing into and out of the cortical surface correspond to anodal and cathodal stimulation [10,11]. However, a preferential direction might not always be clearly defined, such as in the case of subcortical targets. Instead, optimizing the electric field strength might be preferred in this case. This problem was tackled in two prior studies that proposed methods to maximize the field strength in a single target without control of the focality of the resulting field [4,5].

Here, we introduce a novel TES optimization algorithm which controls the field strength in one or more targets while minimizing it elsewhere and at the same time complying with safety constraints and limiting the number of active electrodes. Controlling the target field strength to reach a desired value instead of maximizing it allows for leveraging the trade-off between strength and focality, and for the balanced stimulation of two or more targets. We show that our approach outperforms naïve brute-force search and demonstrate that it succeeds in optimizing montages for the balanced stimulation of the bilateral amygdala.

## 2. Methods

### 2.1 Head Model

We used the example head model from SimNIBS 3.1 (www.simnibs.org) [12] *Ernie* with six tissues compartments, White Matter (WM), Gray Matter (GM), Cerebrospinal Fluid (CSF), Skull, Scalp, and Eyes (figure 1(a)). The standard tissue conductivities in SimNIBS were used [13]. Details of the MR image types and parameters and of the methods used to create the head model are given in [14].

**Figure 1.**
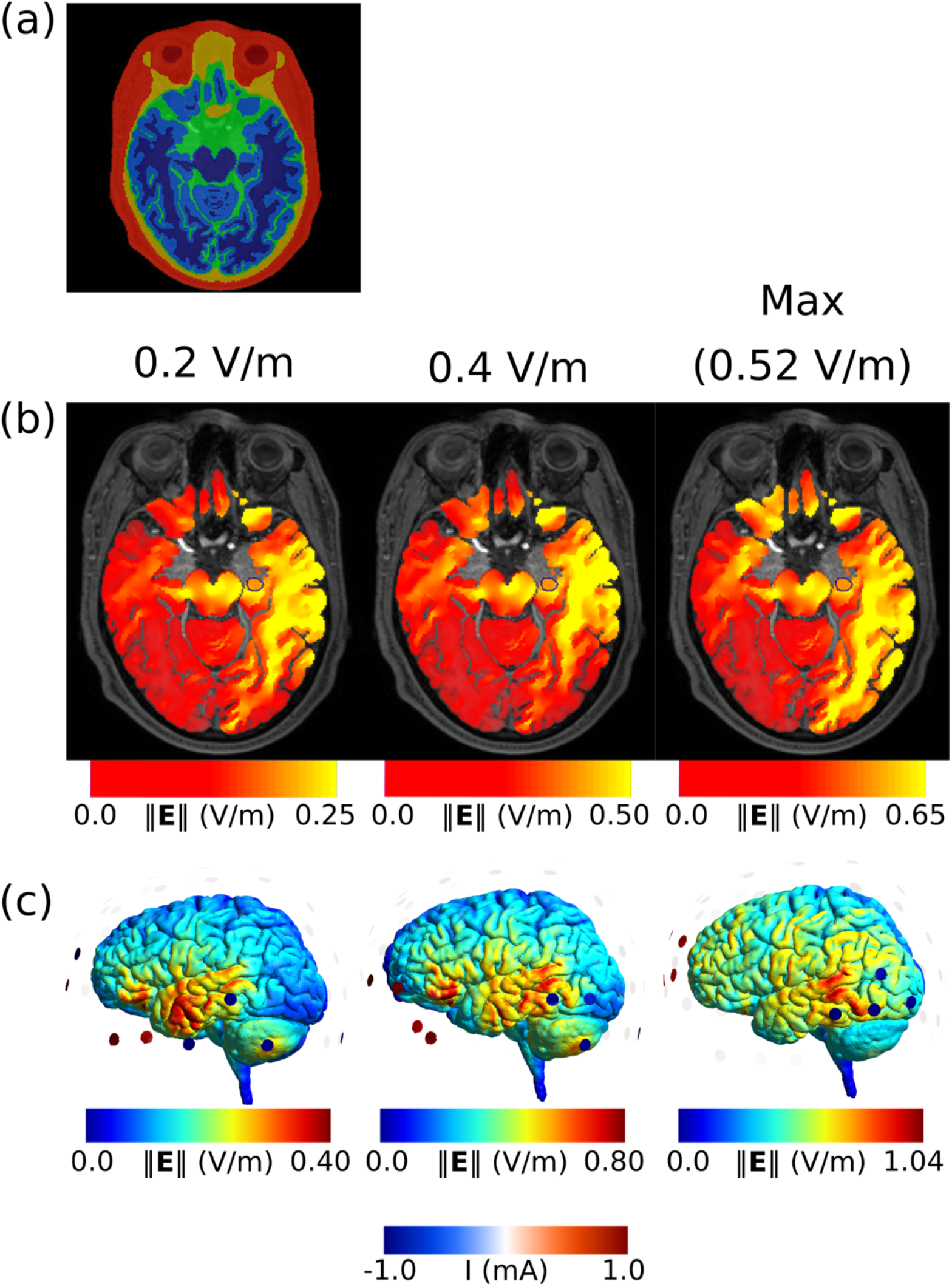
(a) Horizontal slice through the Ernie head model, automatically created by the SimNIBS *headreco* pipeline. The six tissues are white matter (dark blue), gray matter (light blue), CSF (green), skull (yellow) and scalp (red). (b) Optimized electric fields in gray and white matter while controlling the field strength in the left caudal amygdala to be 0.2 V/m, 0.4 V/m, or while maximizing it. The target is delineated in blue. Notice that the color scale changes proportionally with the electric field in the target. (c) Optimized electric fields in the central gray matter surface together with the active electrodes.

### 2.2 Electric Field Simulations

The optimization algorithm builds upon a leadfield matrix ***A*** [9,15], which is constructed by selecting a fixed return electrode and then injecting a unity current sequentially through each remaining electrode. The leadfield allows for quick evaluations of the electric fields produced by any current combination by leveraging the linearity of the field with respect to the injected currents. Here, we used *n* = 74 electrodes placed according to the EEG 10-10 system and performed simulations in SimNIBS 3.1 using Neumann boundary conditions in the electrode surfaces and the MKL PARDISO solver [16].

### 2.3 Mathematical Formulation

The mean electric field strength in a target region Ω_*t*_ is given by

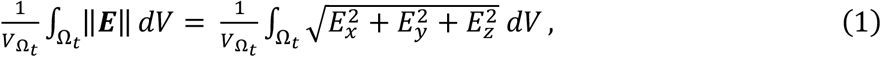

where ***E*** is the electric field and *V*_Ω_*t*__ the volume of the target region. We can approximate the mean of the norm by

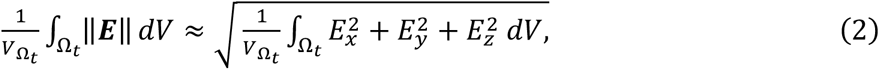

which is a good approximation for a small target region, or if the electric field is approximately constant inside Ω_*t*_. Using this approximation and discretizing the system, we can write the equation above as

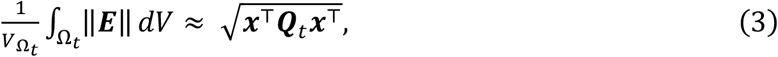

where ***x*** is an *n* × 1 vector of electrode currents, and ***Q**_t_* is a *n* × *n* symmetric positive semidefinite matrix given by

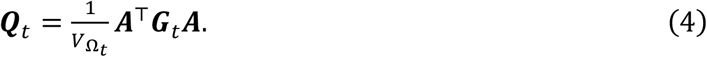

***G**_t_* is a diagonal matrix with zeros in entries outside the target region and element volume values for entries inside the target region.

### 2.4 Optimization Problems

We set up an optimization problem to minimize the field outside the target region, while keeping the mean field strength in the target region at a desired value t, in line with [9]:

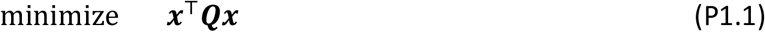

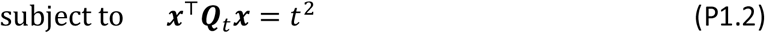

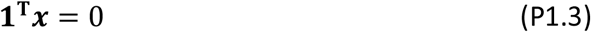

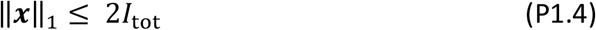

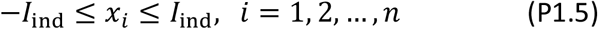

#### Problem 1

Minimize field energy while controlling the electric field strength in the target region.

Here, P1.1 calculates the total field energy, P1.2 controls the field strength in the target, P1.3 enforces Kirchhoff’s law, P1.4 limits the total current injected, and P1.5 limits the current injected through each electrode. Please see [9] for more details. In the following, we only consider the field energy in GM for optimization (P1.1). However, in general, any region can be used.

We can extend Problem 1 to limit the number of active electrodes to *N*, and to control the electric field strength in *n_*t*_* target regions:

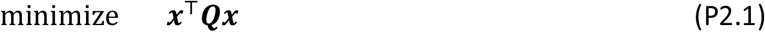

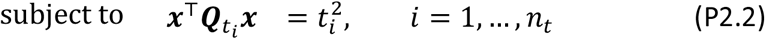

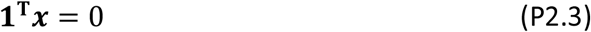

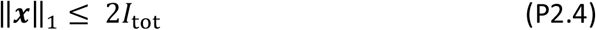

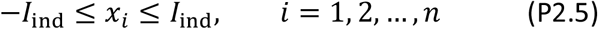

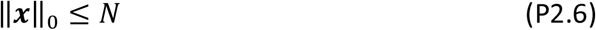

#### Problem 2

Minimize field energy while controlling the electric field strength in *n_*t*_* regions and limiting the number of active electrodes to *N*

### 2.5 Implementation

In contrast to the control of a field component along a specific direction, constraint P1.2 makes problem 1 non-convex. However, this constraint can be dealt with effectively using convex-concave programming (CCP) [17]. This class of optimization algorithms works in problems where each term can be written as a difference of a convex and a concave function, called difference of convex (DC) programming problems. It then proceeds by linearizing the concave part of the functions, thereby obtaining convex optimization problems which can be readily solved.

In order to apply the CCP algorithm, we first substitute the equality constraint P1.2 with an inequality constraint

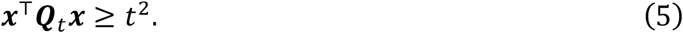

Any solution to the modified problem is also a solution to the original one, as for any point ***x′*** in the feasible region of the modified problem there exists another point 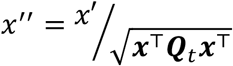, which fulfills both constraint P1.2 and Inequality 5 and has a smaller objective value. Linearizing inequality 5 around a point ***x**_k_*, we obtain the term

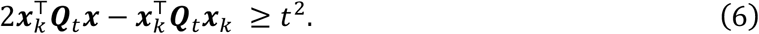

Because of the convexity of the quadratic term, any point that obeys inequality 6 also obeys Inequality 5. We then solve at each step *k* an optimization problem:

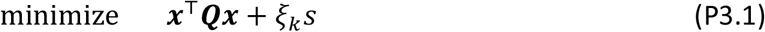

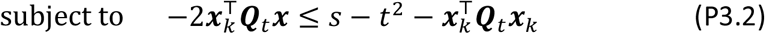

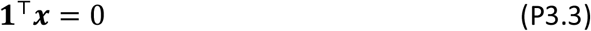

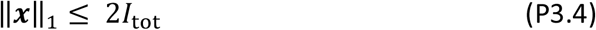

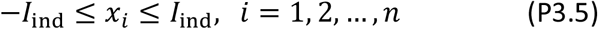

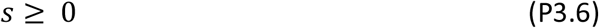

#### Problem 2

Linearization of problem 1 around a point *x_k_*

where *ξ_k_* is a nondecreasing penalty term and *s* is a slack variable. The introduction of the slack variable allows for the algorithm to be initialized at a point ***x***_0_ which violates inequality 5, and for it to better explore the optimization domain and find regions of lower objective value [17]. In order to obtain a feasible solution, we increase the penalty term at each iteration by a factor *μ* > 1, until a maximum value of *ξ*_max_ is reached

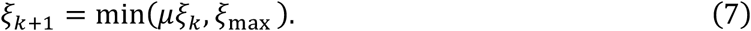

For the current work, we used

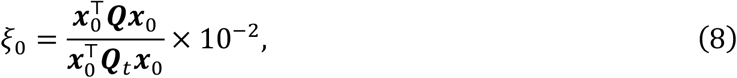

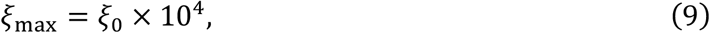

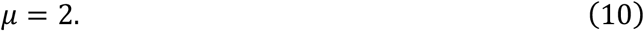

The initial points ***x***_0_ are obtained by solving the constrained eigenvalue problem

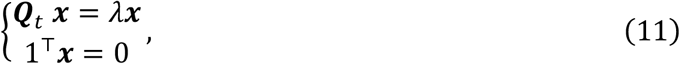

and then scaling the eigenvalues so that they obey constraints (P1.4) and (P1.5). In order to obtain solutions closer to the global optimum, we performed a total of 20 starts for each optimization. We considered that the optimization converged when the objective stopped decreasing and the electric field strength in the target stopped increasing

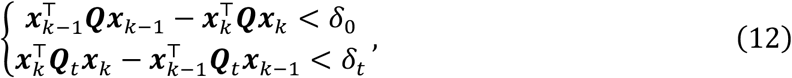

where

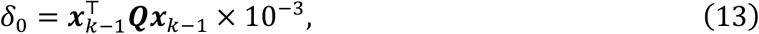

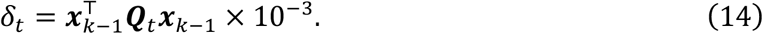

Notice that, if the target strength value *t* is too large, it might not be reachable, which makes problem 1 infeasible. In this case, the slack term *ξ_k_s* dominates and the optimization algorithm will naturally convert to a problem of maximizing field strength in the target region.

In order to deal with many target regions simultaneously (Constraint P2.2), we go through the same steps as with a single constraint, but modify the constrained eigenproblem used for initialization (Equation 11) to use the sum of the ***Q**_t_i__* matrices.

Constraining the number of active electrodes (Constraint P2.6) leads to a combinatorial problem. However, this problem can be efficiently solved using the branch-and-bound algorithm described in [9].

### 2.6 Validation

In order validate our optimization algorithm, we used it to optimize 1000 randomly chosen GM target regions, each with a 10mm radius. To test whether the results obtained were indeed optimal, we compared the solutions to the best value obtained by optimizing the electric fields while controlling directional components (i.e. solving Problem 3 in [9]) in 25 directions equally spaced in a half-sphere.

Afterwards, we performed optimizations while controlling the field strength in two targets simultaneously. Two hundred pairs of target regions were randomly selected in GM. To validate the algorithm, we compared the results with the best values obtained from minimizing the total field energy while controlling the field component in each target independently (Problem 8 in [9]). We searched all combinations of 12 equally spaced directions in a half sphere for one of the targets and 25 directions in a full sphere for the other target, which gives a total of 300 directions. In all cases, we limited the total current injected to 2 mA and the current injected per electrode to 1mA. The target intensity was set to 0.2 V/m. Performance of the optimization was assessed using ratio of the energy (P1.1) of the solutions obtained with the optimization and the search method (ratios <1 indicate better performance of the optimization method).

### 2.7 Subcortical Target

To illustrate the effect of the target field strength on the electric field, we optimized electrode montages while controlling the field strength either in the left amygdala (figure 1(b)-(c)) or the bilateral amygdala (figure 2), and also maximized the field strength in these regions. The current flow through each electrodes was limited to 1 mA, the total current injected to 4 mA and the number of active electrode was limited to 8 using the branch-and-bound algorithm described in [9]. The target intensity was set to 0.2 V/m, 0.4 V/m or maximized. In the two-target case, the later was done while keeping the electric field strength in both targets the same. Even though the images show the electric field in grey and white matter, only the electric field in gray matter was considered during optimization.

**Figure 2.**
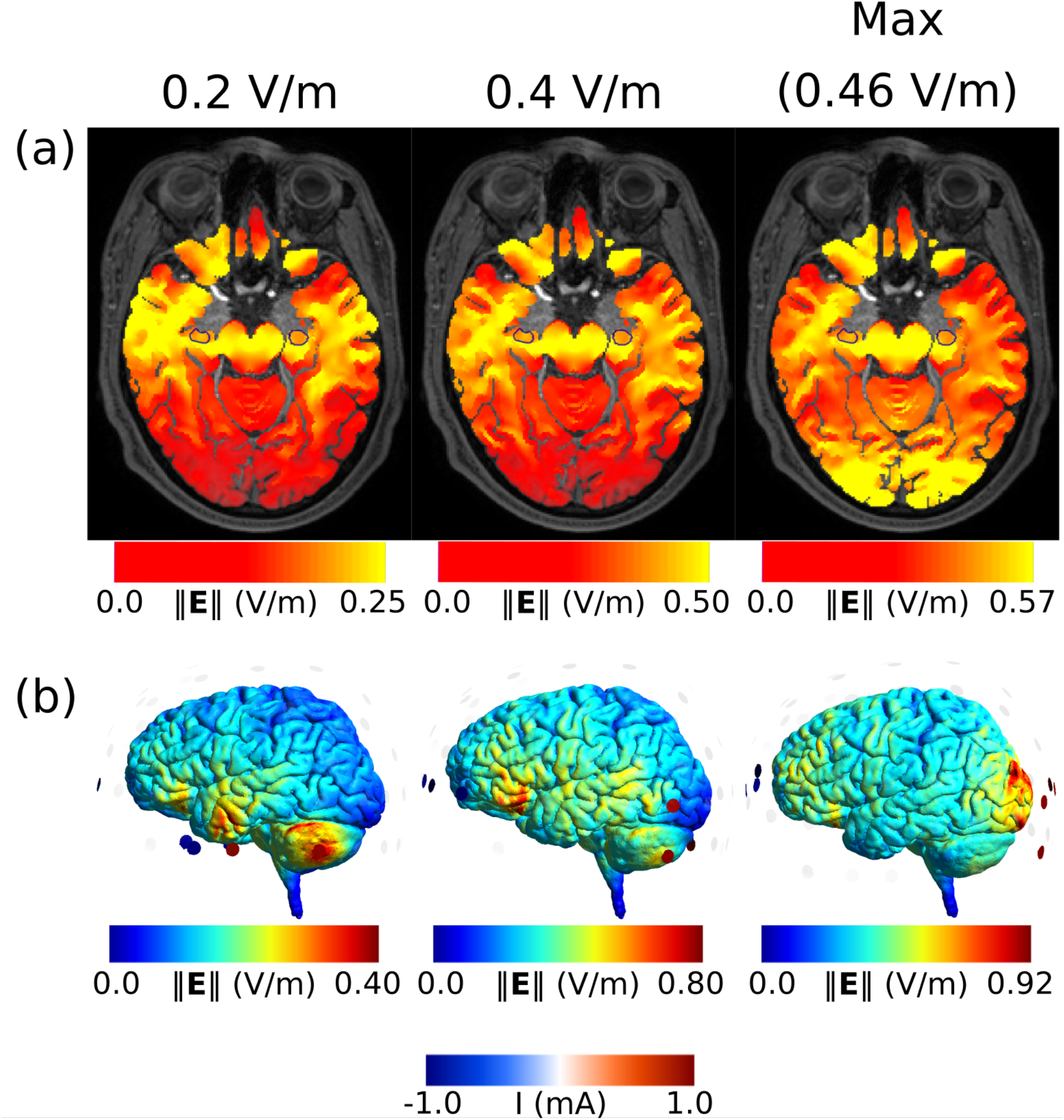
(a) Optimized electric fields in gray and white matter while controlling the field strength inside the two targets delineated in blue. The electric field strength in both targets was set to 0.2 V/m, 0.4 V/m, and increased further until the achieved field strength started differing between both targets. The color scale changes proportionally with the electric field strength in the target. (b) Optimized electric fields shown in the central gray matter surface with the electrode montages overlaid.

## 3. Results

### 3.1 Validation

When controlling the field strength of one target, our optimization approach outperformed the naïve search in all the cases (range of energy ratios between optimized vs searched solutions: 0.87 to 0.99; 95% confidence interval, CI), while running ~1.7 times faster (2.2 vs. 3.7 seconds). For the two-target control, the optimization was better in 99.5% of the cases (range of energy ratios between optimized vs naïve search: 0.82 to 0.97; 95% CI), while running ~3.5 times faster (4.0 vs. 14.5 seconds).

### 3.2 Subcortical Target

The peak electric fields are not located in the subcortical target, but in more superficial structures (figure 1(b)), which is expected from earlier findings [9]. Interestingly, however, as the field strength in the target increases, the fields in superficial regions get disproportionally stronger (i.e. the focality decreases strongly). This is expected due to physical limitations in the distribution of electric fields [3,9]: Increasing the field strength in the target is achieved by increasing the distance between anodes and cathode (Figure 1C), which lowers focality. This suggests that controlling the field strength rather than simply maximizing is preferred to maintain a better focality.

Figure 2 shows the same intensity-focality trade-off for the two target case. Interestingly, the trade-off between 0.2 V/m and 0.4 V/m seems small. The electrode montages changes from being symmetric along the sagittal plane to a frontal-posterior montage as the target intensity increases beyond 0.4 V/m (Figure 2B).

## 4. Discussion

Our new algorithm is capable of optimizing the focality of multichannel TES montages while controlling the field strength in multiple targets. The algorithm performs better than simple search both in terms of the optimality of the solution and time. When compared to the results obtained by maximizing the field strength in the target, the algorithm can strongly improve the focality of the stimulation at merely moderately weaker fields in the target.

Exemplary optimization of the field strength in the uni- and bilateral amygdala reveal that relatively high field strengths can be obtained also for bilateral targeting, but also shows an expected intensity-focality trade-off and stronger fields in cortical regions. Controlling the field strengths in the bilateral amygdala rather than merely maximizing it achieves a balanced montage in which both targets are similarly stimulated.

For superficial targets, it might often be more desirable to control a specific field component in order to ensure that the field is oriented perpendicularly to the cortical surface in the target area. For subcortical targets, however, a preferential direction might not be easily defined and the control of the field strength might be preferred. The new algorithm is computationally efficient and obeys safety and practical constraints, rendering it suited for use in empirical TES studies. It will be released as open source in a future version of the transcranial brain stimulation simulation and optimization software SimNIBS [12].

## 5. Acknowledgments

This study was supported by the Lundbeck foundation (R244-2017-196 and R186-2015-2138), the Novonordisk foundation (NNF14OC0011413) and the NIH (1RF1MH117428-01A1).

## References

[1] Liu A, Vöröslakos M, Kronberg G, Henin S, Krause M R, Huang Y, Opitz A, Mehta A, Pack C C, Krekelberg B, Berényi A, Parra L C, Melloni L, Devinsky O and Buzsáki G 2018 Immediate neurophysiological effects of transcranial electrical stimulation Nat. Commun. 9

[2] Opitz A, Paulus W, Will A and Thielscher A 2015 Anatomical determinants of the electric field during transcranial direct current stimulation Neuroimage 109 2

[3] Dmochowski J P, Datta A, Bikson M, Su Y and Parra L C 2011 Optimized multi-electrode stimulation increases focality and intensity at target J. Neural Eng. 8 046011

[4] Park J H, Hong S B, Kim D W, Suh M and Im C H 2011 A novel array-type transcranial direct current stimulation (tDCS) system for accurate focusing on targeted brain areas IEEE Trans. Magn. 47 882–5

[5] Dmochowski J P, Datta A, Huang Y, Richardson J D, Bikson M, Fridriksson J and Parra L C 2013 Targeted transcranial direct current stimulation for rehabilitation after stroke Neuroimage 75 12–9

[6] Ruffini G, Fox M D, Ripolles O, Miranda P C and Pascual-Leone A 2014 Optimization of multifocal transcranial current stimulation for weighted cortical pattern targeting from realistic modeling of electric fields Neuroimage 89 216–25

[7] Wagner S, Burger M and Wolters C H 2016 An Optimization Approach for Well-Targeted Transcranial Direct Current Stimulation SIAM J. Appl. Math. 76 2154–74

[8] Guler S, Dannhauer M, Erem B, Macleod R, Tucker D, Turovets S, Luu P, Erdogmus D and Brooks D H 2016 Optimization of focality and direction in dense electrode array transcranial direct current stimulation (tDCS) J. Neural Eng. 13 1–14

[9] Saturnino G B, Siebner H R, Thielscher A and Madsen K H 2019 Accessibility of cortical regions to focal TES: Dependence on spatial position, safety, and practical constraints Neuroimage 203 116183

[10] Bindman L J, Lippold O C J and Redfearn J W T 1964 The action of brief polarizing currents on the cerebral cortex of the rat (1) during current flow and (2) in the production of long-lasting after-effects J. Physiol. 172 369–82

[11] Stagg C J and Nitsche M A 2011 Physiological Basis of Transcranial Direct Current Stimulation Neurosci. 17 37–53

[12] Thielscher A, Antunes A and Saturnino G B 2015 Field modeling for transcranial magnetic stimulation: A useful tool to understand the physiological effects of TMS? 2015 37th Annual International Conference of the IEEE Engineering in Medicine and Biology Society (EMBC) (IEEE) pp 222–5

[13] Saturnino G B, Madsen K H and Thielscher A 2019 Electric field simulations for transcranial brain stimulation using FEM: An efficient implementation and error analysis J. Neural Eng. 16

[14] Nielsen J D, Madsen K H, Puonti O, Siebner H R, Bauer C, Madsen C G, Saturnino G B and Thielscher A 2018 Automatic skull segmentation from MR images for realistic volume conductor models of the head: Assessment of the state-of-the-art Neuroimage 174 587–98

[15] Dmochowski J P, Koessler L, Norcia A M, Bikson M and Parra L C 2017 Optimal use of EEG recordings to target active brain areas with transcranial electrical stimulation Neuroimage 157 69–80

[16] Intel 2019 Intel MKL PARDISO - Parallel Direct Sparse Solver Interface

[17] Lipp T and Boyd S 2016 Variations and extension of the convex–concave procedure Optim. Eng. 17 263–87

